# Acute noise causes down-regulation of ECM protein expression in guinea pig cochlea

**DOI:** 10.1101/2020.09.27.315952

**Authors:** Min Shi, Lei Shi, Daxiong Ding, Yiyong Hu, Guowei Qi, Li Zhan, Yuhua Zhu, Ping Lv, Ning Yu

## Abstract

**Objective:** To study the differences in cochlea protein expression before and after noise exposure using proteomics to reveal the pathological mechanism of noise-induced hearing loss (NIHL).

**Methods:** A guinea pig NIHL model was established to test the ABR thresholds before and after noise exposure. The proteomics technology was used to study the mechanism of differential protein expression in the cochlea by noise stimulation.

**Results:** The average hearing threshold of guinea pigs on the first day after noise exposure was 57.00±6.78dB SPL, high-frequency hearing loss was more severe than low frequency; the average hearing threshold on the seventh day after noise exposure was 45.83±6.07dB SPL, and the 4k Hz hearing threshold has the best recovery. The proteomics technology identified 3122 different inner ear proteins, of which six proteins related to the hearing were down-regulation: TenascinC, Collagen type XI alpha two chains, Collagen type II alpha one chain, Thrombospondin 2, Collagen type XI alpha one chain and Ribosomal protein L38, and are enriched in protein absorption, focal adhesion, and extracellular matrix receptor pathways.

**Conclusion:** Impulse noise can affect the expression of differential proteins through focal adhesion pathways. This data can provide an experimental basis for the research on the prevention and treatment of NIHL.

## Introduction

NIHL is a major occupational disease worldwide, with more than 1.6 million new cases every year [1] and is the second most common sensorineural hearing impairment after presbycusis [2, 3]. Li Xingqi [4] pointed out that the hearing loss caused by strong impulse sound stimulation is mainly mechanical damage, while the hearing loss caused by steady-state noise is mainly metabolic damage, indicating that the mechanism of NIHL is closely related to the type of noise exposed. However, there is still a lack of clear experimental evidence at the protein level. The emergence of proteomics technology provides a powerful tool to reveal the molecular mechanism of noise-induced hearing loss at the protein level. TMT (Tandem Mass Tags) technology is a quantitative proteomics in recent years. The widely used high-throughput screening technology can be used to label the N-terminus and amino acid side chain groups of protein peptides and realize the quantification of proteins between different samples by mass spectrometry. In this study, TMT technology was used to study the difference of protein expression in the cochlea of guinea pigs before and after noise exposure, and biological analysis was carried out to reveal the marker proteins, potential pathogenesis and intervention targets after noise-induced hearing loss.

## Materials and methods

### Animals

Eight healthy adult male Dunkin Hartley guinea pigs, weighing 250-300g, and sensitive to auricle reflex, were purchased from Beijing Keyu Animal Breeding Center. All animals had no history of noise exposure, had never used ototoxic drugs, and had normal tympanic membranes by otoscope. The experimental animals were randomly divided into three groups, namely the normal control group (n=2), the first day group after noise exposure (n=3) and the seventh day group after noise exposure (n=3). All experimental operations were approved by the Animal Ethics Committee of the Chinese People’s Liberation Army General Hospital.

### Impulse noise exposure

Place 6 awake guinea pigs in a special metal cage, fix the head of the guinea pig in the center, and ensure that its head does not rotate during noise exposure, and ensure that both ears of the guinea pig receive the same impulse noise stimulation. Place the special impulse noise excitation device at the front level of the animal’s head, 4-5cm away from the front edge of the guinea pig ears, and give 20 consecutive pressure peaks of 165 dB SPL, pulse width of 0.25ms, and interval of 6.5s Impulse noise lasts for two days, a total of 40 times.

### Auditory brainstem response, ABR

Anesthetize the animal by intraperitoneal injection of 1% sodium pentobarbital (0.4ml/g) and place the animal on an insulation blanket after breathing and heartbeat are stable. Audiometry is conducted in a sound-proof shielded room, using American TDT audiometry equipment and Biosig audiometry analysis software. Connect the TDT electrode: the recording electrode is inserted into the skull parietal subcutaneously from the midpoint of the upper edge of the animal’s double auricles and the junction of the skull, the reference electrode is inserted under the skin of the test ear, and the ground electrode is inserted under the skin of the opposite ear to ensure the test resistance Less than 1kΩ, the earphone is placed 0.2cm at the mouth of the ear canal outside the test ear. The stimulus sounds are “Click”, pure tones “tone burst” (4kHz, 8kHz, 16kHz), the filter width is set to 300-3000Hz, and the stimulation duration is 10ms, and the overlay is 1024 times, The maximum stimulus intensity is 90dB SPL, which is reduced by 10dB SPL. When approaching the hearing threshold, the stimulus intensity decreases by 5dB SPL. When there is an irregular and illegible waveform, it increases by 5dB SPL. The lowest stimulus intensity that induces a repeatable regular wave is recorded as the ABR threshold.

### TMT (Tandem Mass Tags) technology

The three groups of animals were quickly sacrificed after anesthesia, decapitated and removed the bilateral cochlea, rinsed with PBS, quickly placed in liquid nitrogen for 10 minutes, and then placed in a −80°C refrigerator.

TMT experiment process: Cochlear sample preparation (extracted protein) → SDS-PAGE electrophoresis to detect protein quality → (sample protein qualified) Trypsin digestion → TMT reagent labeling and classification → mass spectrometry analysis → software analysis and statistical analysis → bioinformatics analysis (GO analysis, KEGG Pathway)

Grind the guinea pig cochlea with a mortar. For protein extraction and quantification methods, please refer to previous literature reports [5]. Draw a protein quantitative standard curve based on the standard product and get the correlation coefficient R^2^=0.9856>0.95. It is believed that there is a linear relationship between the amount of protein and the OD absorbance. 10% SDS-PAGE electrophoresis determined that the total protein was effectively separated in the molecular weight range of 15-220 kDa, and the protein bands in the three sample groups had similar band patterns.

The experimental methods of trypsin digestion of protein samples and peptide labeling with TMT reagent can refer to previous literature reports [5]. Sample peptides of each group were taken about 100μg and labeled according to the instructions of the TMT kit. The normal control group, the first day group and the seventh day group were subjected to 3 biological replicates.

Finally, the capillary efficient liquid chromatography and mass spectrometry identification. Each sample was separated by a nanoliter flow rate HPLC liquid phase system, and then analyzed by Orbitrap Fusion Lumos mass spectrometer.

Raw mass spectrometry data: The raw mass spectrometry data is a RAW format file, and the software Proteome Discoverer is used for qualitative and quantitative analysis.

Database: uniprot-Cavia+porcellus

Search the database: upload the RAW file to Proteome Discoverer when searching the database, select the established database, and then search the database.

All experimental data are expressed as mean±standard deviation (x±s), and the experimental data are analyzed using SPSS 20.0 statistical software package. The differences between the three groups were compared by one-way analysis of variance. A *p*-value less than 0.05 was considered statistically significant.

## Results

### Noise exposure causes about 20 dB of hearing loss

In order to exclude guinea pigs with poor hearing or deafness, all guinea pigs received ABR test before noise exposure; the ABR threshold test results of normal control group, first day after noise exposure, and seventh days after noise exposure are shown in Table 1. The hearing test results showed that the ABR threshold of normal guinea pigs was 26.88±8.08dB SPL, and the average hearing threshold increased to 57.00±6.78dB SPL within first day after noise exposure. The damage was the most serious at 16k Hz, and the high-frequency hearing loss was more serious than the low-frequency. The seventh day average hearing threshold can be recovered to 45.83±6.07dB SPL, the recovery is best at 4k Hz, and above 4k Hz is poor.

**Table 1.** Three groups of guinea pig ABR hearing results (dB SPL, x±s)

| group | click | 4k | 8k | 16k |
| --- | --- | --- | --- | --- |
| Pre | 26.88±8.08 | 10.94±2.63 | 12.81±4.32 | 25.94±9.72 |
| NE1d | 57.00±6.78 | 61.00±11.58 | 64.00±9.70 | 66.00±12.81 |
| NE7d | 45.83±6.07 | 35.00±11.90 | 58.33±12.47 | 58.33±16.75 |
| F | 32.96 | 75.31 | 101.96 | 23.51 |
| P | P<0.01 | P<0.01 | P<0.01 | P<0.01 |

Annotation: Pre represents the normal control group; NE1d presents first day after noise exposure group; NE7d presents seventh days after noise exposure group.

### TMT technology screened out about 90 different proteins

Through cluster analysis of expression patterns among multiple samples, the results show that the repeatability between samples is good. After screening with Peptide FDR≤0.01, 3122 proteins were identified. Compared with the normal control group, the protein expression of the group first day after noise exposure was mainly up-regulated, and the protein expression of the group seventh days after noise exposure was mainly down-regulated compared with the group first day after noise exposure and the normal control group(Figure 1).

**Figure 1.**
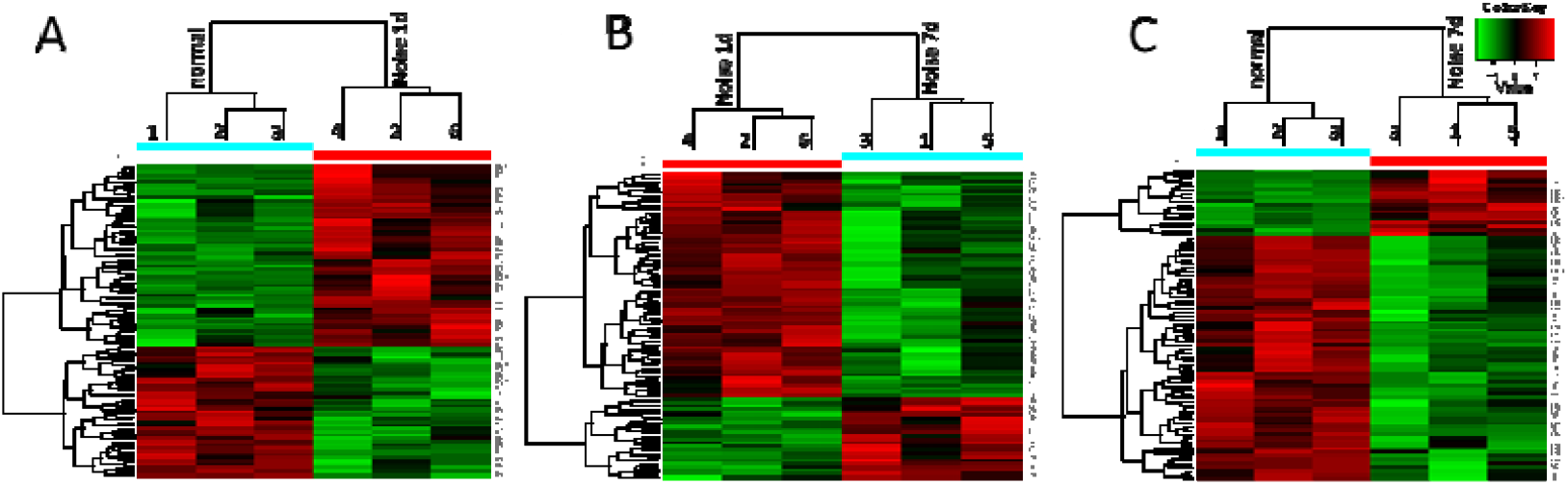
The cluster analysis of expression patterns among multiple samples showed that the expression of inner ear protein was significantly different at different time points after noise exposure. A: First day after noise exposure, the protein expression of the group is mainly up-regulated relative to the normal control group. B: The protein expression of the seventh days after noise exposure group was mainly down-regulation compared with the first day after noise exposure group. C: The protein expression in the group seventh days after noise exposure was mainly down-regulation compared with the normal control group. A total of 3122 proteins were identified.

Use Proteome Discoverer proteomics analysis software to realize TMT quantification. According to the quantitative results, set at 1.2 fold change (FC) and P<0.05 threshold conditions (FC>1.2 and P<0.05 for up-regulation, FC≤0.833 and P<0.05 for down-regulation, 0.833<FC<1.2 or p>0.05, it is considered that there is no significant change in expression) Differential protein screening, the results show that: first day after noise exposure, there are 96 kinds of proteins that are significantly differently expressed than the normal group, of which 56 are up-regulated and 40 are down-regulated;A total of 90 proteins were significantly differently expressed in the seventh days after noise exposure group than the first day after noise exposure group, of which 24 were up-regulated and 66 were down-regulated;seventh days after noise exposure, there are 84 kinds of proteins that are significantly differently expressed than the normal group, of which 18 are up-regulated and 66 are down-regulated; see Figure 2 volcano graph and histogram.

**Figure 2.**
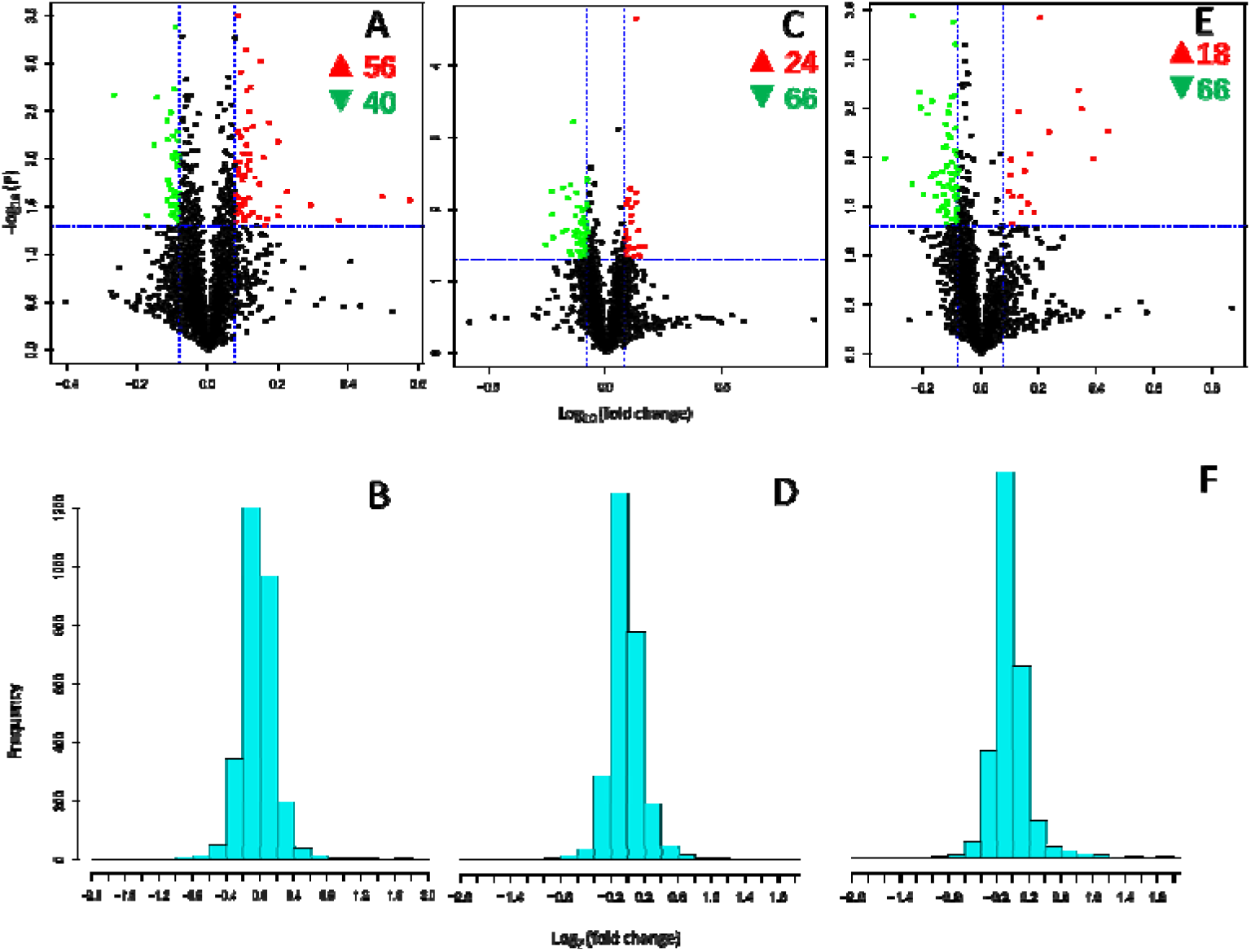
Noise exposure significantly affects cochlear protein expression. In the volcano map, the abscissa is the logarithmic transformation of the multiple of the difference, and the ordinate is the logarithm transformation of the significant difference P-value (P<0.05 is statistically significant); the green and red dots indicate significant differences Protein (FC≤0.833-green, down-regulated; FC>1.2-red, up-regulated), the black dots are proteins with no significant changes. The ordinate in the histogram is the protein quantity, and the abscissa is the Log2 logarithm of the quantitative ratio FC of the two sets of samples; Among them, the relative quantitative ratio of most proteins in each two groups of samples is close to 1, and the distribution is more concentrated. The farther the distance is from 1, the difference is obvious. A, B. first day after noise exposure group and normal control group, a total of 96 proteins with significant differential expression, of which 56 are up-regulated and 40 are down-regulated; C, D. seventh days after noise exposure and first day after noise exposure Group, there are a total of 90 proteins with significant differential expression, of which 24 are up-regulated and 66 are down-regulated; E, F: seventh days after noise exposure group and normal control group, a total of 84 proteins with significant differential expression, of which 18 are up-regulated and 66 are down-regulated.

Analysis by Gene Ontology (GO) found that the expression of 6 proteins related to hearing was significantly down-regulation

About 90 differential proteins screened out by the TMT technology were then subjected to GO function enrichment analysis. In the comparison between the seventh days after noise exposure group and the first day after noise exposure group, it was found that the expression of 6 hearing-related proteins was significantly down-regulated : Tenascin C (TNC); Collagen type XI alpha 2 chain (COL11A2); Collagen type II alpha 1 chain (COL2A1); Thrombospondin 2 (THBS2); Collagen type XI alpha 1 chain (COL11A1); Ribosomal protein L38 (RPL38) expression down. Compared with the normal group, the first day after noise exposure group and the seventh days after noise exposure group had up-regulated protein expression, but they were not screened out because the difference was not statistically significant. The expression of 6 differential proteins in the three groups of the cochlea in Figure 3.

**Figure 3.**
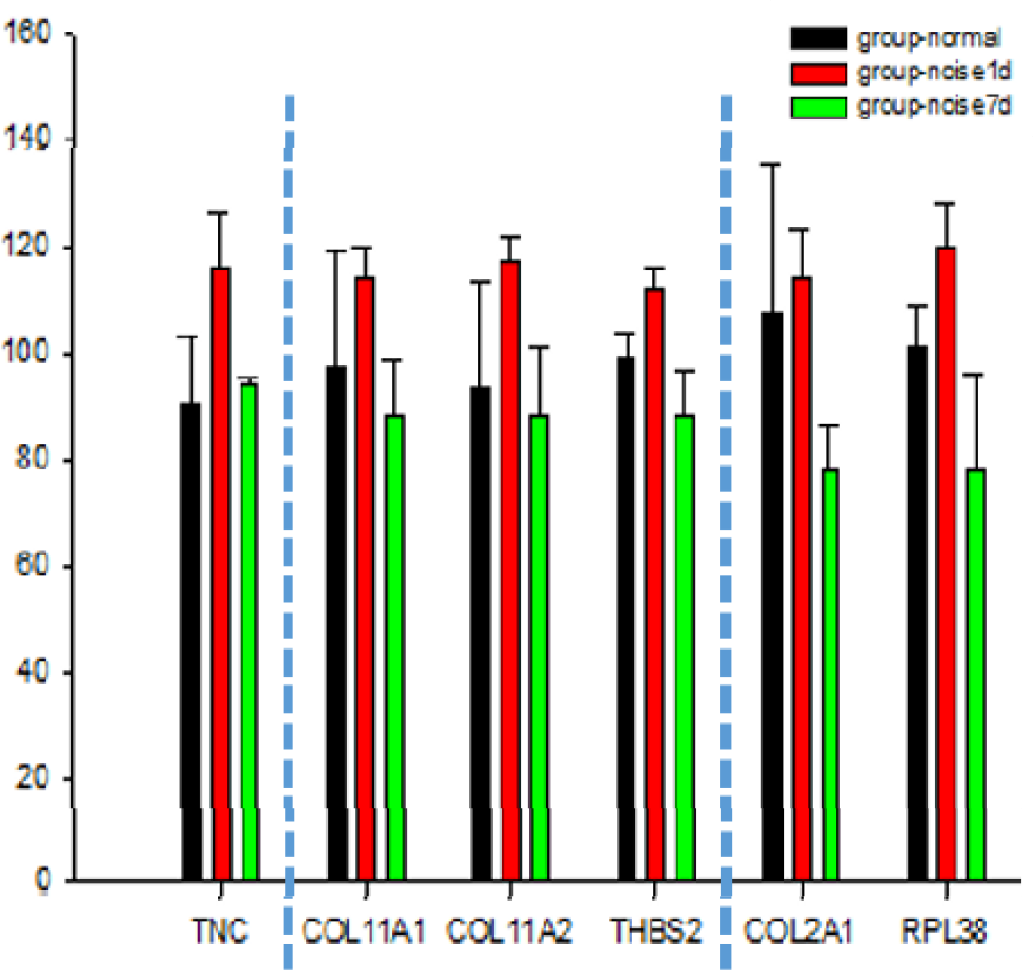
Noise exposure has a significant effect on the expression of 6 proteins including TNC, COL11A1, COL11A2, THBS2, COL2A1 and RPL38. The abscissa represents the 6 differential proteins related to hearing, and the ordinate represents the relative quantitative value of protein markers; the expression of the 6 proteins before and after noise exposure was significantly increased, increasing by 6%-30%; on the seventh day after noise exposure, TNC protein expression was up-regulated by about 3%, while the down-regulation of COL11A1, COL11A2 and THBS2 protein expression was within 10%, and the down-regulation of COL2A1 and RPL38 protein expression was more than 20%. Black represents the normal group; red represents the first day group after noise exposure; green represents the seventh day group after noise exposure. TNC stands for TenascinC; COL11A1 stands for Collagen type XI alpha 1 chain; COL11A2 stands for Collagen type XI alpha 2 chain; COL2A1 stands for Collagen type II alpha 1 chain; RPL38 stands for Ribosomal protein L38; THBS2 stands for Thrombospondin 2.

Protein digestion and absorption, Focal adhesion and ECM-receptor interaction signal pathways are the most important biochemical metabolic pathways and signal transduction pathways for the six different proteins.

Different proteins coordinate with each other to perform their biological behaviors in organisms. Pathway analysis helps to further understand their biological functions. In this study, the main biochemical metabolic pathways and signal transduction pathways involved in protein were determined through KEGG Pathway analysis. The KEGG function annotations of different proteins are shown in Table 2 and Figure 4. Comparing the seventh day group after noise exposure with the first day group after noise exposure, the functional annotations of KEGG Pathway enriched by differential proteins related to hearing include protein digestion and absorption, extracellular matrix receptor interaction (ECM-Receptor interaction), focal adhesion signal. COL11A1, COL11A1, and COL2A1 are all enriched in protein digestion and absorption pathway, COL2A1, THBS2, and TNC proteins are all enriched in ECM-receptor interaction pathway, focal adhesion signal pathway (Table 2).

**Table 2.** Three signal pathways of differential protein enrichment related to deafness

|  | COL11A1 | COL11A2 | COL2A1 | THBS2 | TNC | RPL38 |
| --- | --- | --- | --- | --- | --- | --- |
| Protein digestion and absorption | √ | √ | √ |  |  |  |
| Focal adhesion |  |  | √ | √ | √ |  |
| ECM-receptor interaction |  |  | √ | √ | √ |  |

1. COL11A1, COL11A2, COL2A1 are enriched in the protein digestion and absorption signal pathway; 2. COL2A1, THBS2, and TNC are enriched in Focal adhesion signal pathway; 3. COL2A1, THBS2, and TNC are enriched in the ECM-receptor interaction signal pathway.

**Figure 4.**
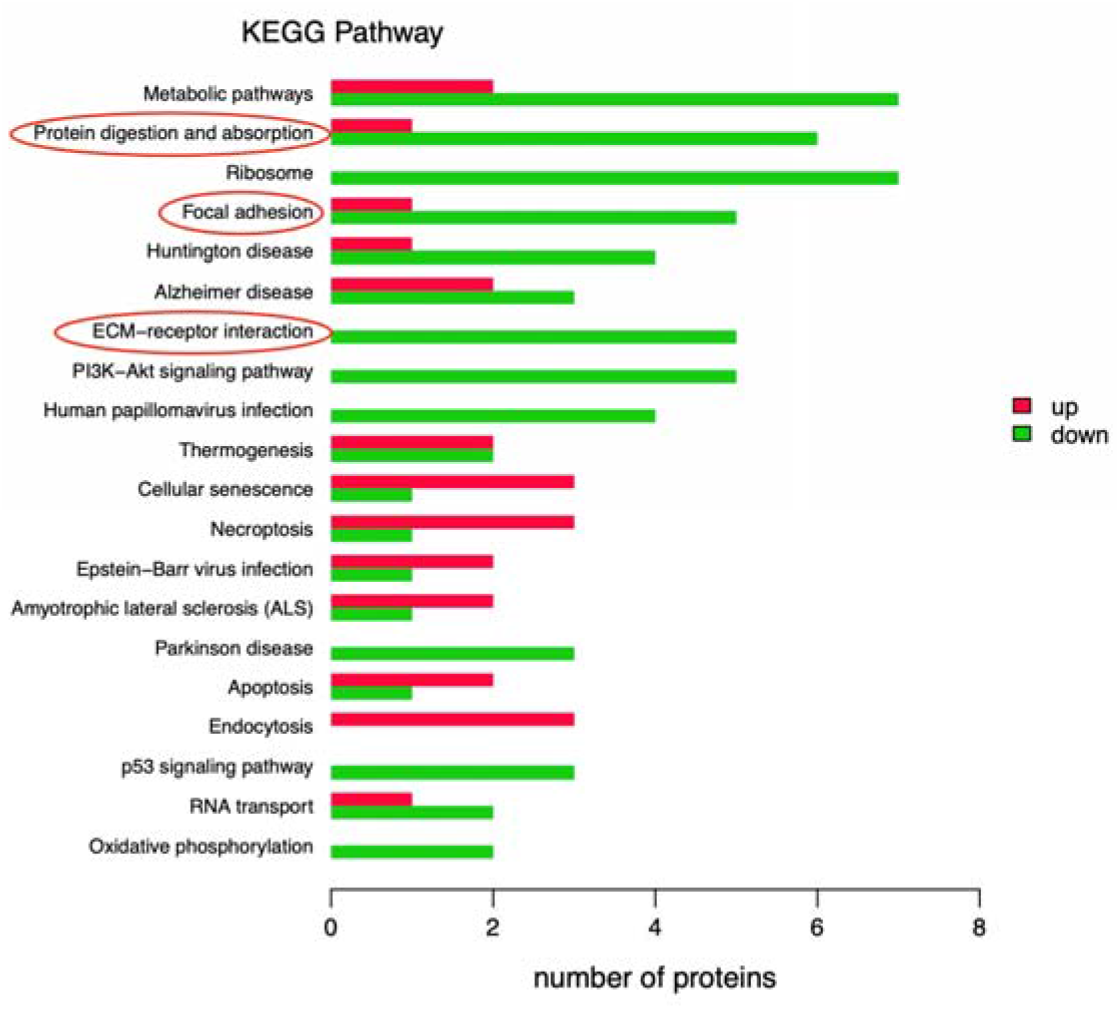
KEGG Pathway analysis found that the three pathways of Protein digestion and absorption, ECM-receptor interaction and Focal adhesion are the most important biochemical metabolic pathways and signal transduction pathways of differential proteins. The abscissa represents the number of differential proteins enriched, and the ordinate represents the description of each KEGG Pathway, that is, the number of differential proteins identified in this experiment under a certain KEGG signaling pathway. Red means up regulation and green means down regulation. Comparing the seventh day post-noise exposure group (NE7d) with the first day post-noise exposure group (NE1d), the functional annotations of KEGG Pathway enriched with differential proteins (related to deafness) include protein digestion and absorption, and extracellular matrix receptors ECM-receptor interaction pathway, focal adhesion signal, the red box marks the functional annotation of the signal pathway enriched by the difference protein (related to deafness) in this experiment.

## Discussion

### Characteristics and parameters of hearing loss in guinea pigs caused by impulse noise exposure

Chen Zhiting [6] found that more than 50 times of 145dB SPL impulse noise exposure can increase the hearing threshold by more than 70dB SPL on average. In this study, the average hearing threshold increased by 20dB SPL after 165dB SPL impulse noise exposure for 40 times. This difference indicates that the increase in impulse noise stimulation Sound intensity, reduce the number of exposures, and less damage to the auditory system. They also found that hearing recovery was the fastest within one week after exposure, which is consistent with the change trend of hearing thresholds in animals after acute noise damage in this study.

### Impulse noise damage affects the proteomics of guinea pig inner ear

In recent years, proteomics related research on NIHL has gradually become a research hotspot. In the past, Samson [7] used antibody microarray technology to quantify the exposure of chinchillas to 112dB SPL noise for 2 hours, and then 1st day after noise exposure and 7th days after noise exposure. 28th days after noise exposure, the changes in protein expression in different regions of the chinchilla cochlea were detected, indicating that the p38/MAPK signal transduction activated by the focal adhesion signaling pathway plays a key role in the mechanism of NIHL.

Yukihide [8] used iTRAQ proteomics method to divide 24 mice exposed to 120dB SPL noise for 2 hours into 4 groups to identify different regulatory proteins of dexamethasone in the mouse cochlea, and the results show that myelin protein and heat shock protein 70 are closely related to noise-induced hearing loss, and dexamethasone significantly regulates the expression level of these proteins in the mouse cochlea.

In this study, the TMT proteomics method was used to identify the difference in protein expression in the cochlea under the influence of noise and screened out 6 kinds of hearing related proteins that were significantly down-regulated, TNC; COL11A2; COL2A1; THBS2; COL11A1; RPL38. Except RPL38, all belong to the ECM family.

There are 3 types of collagen in the 6 different proteins: COL11A2, COL11A1, COL2A1, all of which were down-regulation on the seventh day after noise exposure. Collagen in the ECM protein family is a fibrous structural protein, mainly involved in the construction of cartilage [9, 10]. Previous studies have found that mutations in the COL11A2 gene can cause autosomal recessive non-syndromic sensorineural hearing loss (DFNB53) and autosomal dominant non-syndromic sensorineural hearing loss (DFNA13). The loss of collagen fibers in the mouse tectorial membrane will affect the function of outer hair cells [11-13]; the COL11A1 gene is expressed in the mouse auditory vesicles [14], Booth [15] used splicing variants in the study it was found that COL11A2 gene mutations can cause DFNA37; Cao [16] found that COL2A1 knockout mice have hearing loss and vestibular function reduction, and its mutations cause sensorineural hearing loss in patients with Type 1 Stickler syndrome [17,18]. In the inner ear, type II collagen and type XI collagen are located in the tectorial membrane. It is a colloidal structure composed of extracellular material, located on the reticulum plate of the spiral organ, rich in actin, Type II collagen and type XI collagen. The tectorial membrane plays a vital role in the sensory transmission of sound [19]. Normally, the tips of the three-dimensional cilia on the outer hair cells are embedded in the tectorial membrane, while the tips of the cilia on the inner hair cells are adjacent to this structure. The shearing movement between the apex of the hair cell cilia and the tectorial membrane causes the mechanical displacement of the top cilia of the inner hair cell, thereby opening the mechanically sensitive conduction channel. On the first day after impulse noise stimulation, the expression of COL11A2 and COL11A1 were up-regulated by about 15%, and the expression of COL2A1 was up-regulated by about 6%, that is, the expression of ECM protein increased, which in turn changed the arrangement of collagen fibers on the tectorial membrane to make it from an orderly parallel arrangement to a random arrangement, which changes the mechanical properties of the tectorial membrane and destroys the shearing movement of the cilia and the tectorial membrane at the top of the outer hair cells, resulting in the destruction of hearing [13]. After the recovery period, the hearing threshold is increased compared with before. The down-regulated of COL11A2 and COL11A1 is within 10%, and the down-regulated of COL2A1 is more than 20%, and the expression of ECM protein decreases rapidly. It is speculated that the arrangement of collagen fibers on the tectorial membrane has some recovery. It shows that ECM protein plays a vital role in hearing loss and repair during noise exposure.

Impulse noise activates multiple signal transduction pathways in the inner ear of guinea pigs

Pathway significant enrichment method is the same as GO functional enrichment analysis. Through Pathway significant enrichment, the most important biochemical metabolic pathways and signal transduction pathways involved in differentially expressed proteins can be determined. In the data analysis, we screened significant pathways and found that down-regulated differential proteins (ECM proteins) are enriched in protein digestion and absorption pathways, and ECM-receptor interaction pathway, focal adhesion signal pathway (Figure 4). Focal adhesion is a special structure formed by the contact point between cells and extracellular matrix. It is mainly involved in the structural connection between membrane receptors and the actin cytoskeleton. Signal molecules initiate downstream signal transduction events, which ultimately lead to the actin cytoskeleton. The recombination, cell shape and motility, and gene expression changes, the down-regulated expression of this pathway can attenuate the activation of the p38/MAPK signaling pathway [20]. first day after noise exposure, the animal’s hearing threshold dropped rapidly, and the expression of ECM protein was up-regulation. At the same time, oxidative stress in the inner ear up-regulated Fas ligand, which combined with the corresponding receptor would lead to the formation of a death-inducing signal complex [21], so noise-induced up-regulation of Fas signals may be an important upstream event in NIHL, because these receptors mediate the signal cascade, and finally lead to apoptosis through p38/MAPK signaling [22]; after a seven day recovery period, ECM protein expression is down-regulated, the corresponding oxidative stress is weakened, and the apoptosis of inner ear cells is reduced. Currently, p38/MAPK inhibitors have been shown to protect the inner ear from noise damage [23]. Interestingly, TNC protein is related to hearing loss [24,25]. Studies have found that it is a new pathogenic gene of DFNA13 [26], but it also has repair effect. It was found that its down-regulated expression was detected on the seventh day of the study, but it’s increased by about 3% compared to first day.It is very likely that during the acute injury period on the first day after noise exposure, the oxidative stress pathway was activated, which stimulated the repair of TNC protein. The increase in TNC protein exceeded 20%. In the repair period, the corresponding oxidative stress attenuation, reducing the apoptosis of inner ear cells, the repairing effect of TNC protein is weakened, and the expression of TNC protein is gradually weakened during the repairing period, but it is still higher than normal.

It is worth noting that the RPL38 protein is located in the cytoplasmic ribosome and is involved in ribosome assembly, middle ear morphogenesis, sensory perception of sound and other functions. It does not belong to the ECM family, and information related to deafness is not yet known. it was found that its expression was down-regulation in the seventh days after noise exposure group compared with the first day after noise exposure group in this study. In order to explore whether it is a potential biomarker of NIHL, in the next study, the RPL38 protein will be selected. Through qualitative and quantitative verification of the upstream and downstream expression of the protein in the cochlea, it is also necessary to explore the location of the protein in the cochlea.

In general, this proteomics study on noise-induced inner ear damage shows that the expression of ECM protein in the cochlea is different before and after acute noise damage. Bioinformatics analysis suggests that the focal adhesion signaling pathway mediated by ECM protein participates in the initiation of the inner ear cell death process through a cascade reaction, Which can provide new methods and ideas for the research of NIHL prevention and treatment.

## Acknowledgements

This article was funded by the Special Youth Project of the PLA General Hospital (QNC19051), the Active Health Project of the Ministry of Science and Technology (2020YFC2004001) and the Municipal-School Cooperation Project of Nanchong Science and Technology Bureau (19SXHZ0417)

## References

[1] Leigh J, Macaskill P, Kuosma E, et al. Global burden of disease and injury due to occupational factors. Epidemiology. 1999; 10(5): 626–631.

[2] Sliwinska-Kowalska M, Pawelczyk M. Contribution of genetic factors to noise-induced hearing loss: a human studies review. Mutat Res. 2013; 752(1): 61–65.

[3] Stucken EZ, Hong RS. Noise-induced hearing loss: an occupational medicine perspective. Curr Opin Otolaryngol Head Neck Surg. 2014; 22(5): 388–393.

[4] Li Xingqi, Sun Jianhe, Yang Shiming. Pathophysiology of the cochlea. Beijing: People’s Military Medical Press. 2011.

[5] Li Qiang. Effects of four foods on the activity of digestive enzymes in the midgut of Spodoptera exigua and midgut proteomics. Shandong Agricultural University. 2017.

[6] Chen Zhiting, Wang Fangyuan, Ji Fei, et al. Analysis of auditory brainstem response in miniature pigs and changes after impulse noise exposure. Chinese Journal of Otology. 2016; 14(6): 735–739.

[7] Samson J, Hu Bohua, Mohammad HK, et al. Noise induced changes in the expression of p38/MAPK signaling proteins in the sensory epithelium of the inner ear. J Proteomics. 2011; 75(2): 410–424.

[8] Maeda Y, Fukushima K, Kariya S, et al. Dexamethasone Regulates Cochlear Expression of Deafness-associated Proteins Myelin Protein Zero and Heat Shock Protein 70, as Revealed by iTRAQ Proteomics. Otol Neurotol. 2015; 36(7): 1255–1265.

[9] Myllyharju J, Kivirikko KI. Collagens and collagen-related diseases. Ann Med. 2001; 33(1): 7–21.

[10] Keene DR, Oxford JT, Morris NP. Ultrastructural localization of collagen types II, IX, and XI in the growth plate of human rib and fetal bovine epiphyseal cartilage: type XI collagen is restricted to thin fibrils. J Histochem Cytochem. 1995; 43(10): 967–979.

[11] Chakchouk I, Grati M, Bademci G, et al. Novel mutations confirm that COL11A2 is responsible for autosomal recessive non-syndromic hearing loss DFNB53. Mol Genet Genomics. 2015; 290(4): 1327–1334.

[12] Chen W, Kahrizi K, Meyer NC, et al. Mutation of COL11A2 causes autosomal recessive non-syndromic hearing loss at the DFNB53 locus. J Med Genet. 2005; 42(10): e61.

[13] McGuirt WT, Prasad SD, Griffith AJ, et al. Mutations in COL11A2 cause non-syndromic hearing loss (DFNA13). Nat Genet. 1999; 23(4): 413–419.

[14] Yoshioka H, Iyama K, Inoguchi K, et al. Developmental pattern of expression of the mouse alpha 1 (XI) collagen gene (Col11a1). Dev Dyn. 1995; 204(1): 41–47.

[15] Booth KT, Askew JW, Talebizadeh Z, et al. Splice-altering variant in COL11A1 as a cause of nonsyndromic hearing loss DFNA37. Genet Med. 2019; 21(4): 948–954.

[16] Cao C, Oswald AB, Fabella BA, et al. The CaV1.2 L-type calcium channel regulates bone homeostasis in the middle and inner ear. Bone. 2019; 125: 160–168.

[17] Alexander P, Poulson A, McNinch A, et al. Type I membranous anomaly in Stickler syndrome. Ophthalmic Genet. 2018; 39(1): 147.

[18] Leung L, Hyland JC, Young A, et al. A novel mutation in intron 11 of the COL2A1 gene in a patient with type 1 Stickler syndrome. Retina. 2006; 26(1): 106–109.

[19] Richardson GP, Lukashkin AN, Russell IJ, et al. The tectorial membrane: one slice of a complex cochlear sandwich. Curr Opin Otolaryngol Head Neck Surg. 2008; 16(5): 458–464.

[20] Philips A, Roux P, Coulon V, et al. Differential effect of Rac and Cdc42 on p38 kinase activity and cell cycle progression of nonadherent primary mouse fibroblasts. Biol Chem. 2000; 275(8): 5911–5917.

[21] Suzuki M, Aoshiba K, Nagai A. Oxidative stress increases Fas ligand expression in endothelial cells. J Inflamm (Lond). 2006; 3: 11.

[22] Juo P, Kuo CJ, Reynolds SE, et al. Fas activation of the p38 mitogen-activated protein kinase signalling pathway requires ICE/CED-3 family proteases. Mol Cell Biol. 1997; 17(1): 24–35.

[23] Tabuchi K, Oikawa K, Hoshino T, et al. Cochlear protection from acoustic injury by inhibitors of p38 mitogen-activated protein kinase and sequestosome 1 stress protein. Neuroscience. 2010; 166(2): 665–670.

[24] Pillers DA, Kempton JB, Duncan NM, et al. Hearing loss in the laminin-deficient dy mouse model of congenital muscular dystrophy. Mol Genet Metab. 2002; 76(3): 217–224.

[25] Barker DF, Hostikka SL, Zhou J, et al. Identification of mutations in the COL4A5 collagen gene in Alport syndrome. Science. 1990; 248(4960): 1224–1227.

[26] Zhao Y, Zhao F, Zong L, et al. Exome sequencing and linkage analysis identified tenascin-C (TNC) as a novel causative gene in nonsyndromic hearing loss. PLoS One. 2013; 8(7): e69549.

